# Ten simple rules for structuring papers

**DOI:** 10.1101/088278

**Authors:** Brett Mensh, Konrad Kording

## Abstract

Good scientific writing is essential to career development and to the progress of science. A well-structured manuscript allows readers and reviewers to get excited about the subject matter, to understand and verify the paper’s contributions, and to integrate these contributions into a broader context. However, many scientists struggle with producing high-quality manuscripts and typically get little training in paper writing. Focusing on how readers consume information, we present a set of 10 simple rules to help you get across the main idea of your paper. These rules are designed to make your paper more influential and the process of writing more efficient and pleasurable.

## Introduction

Writing and reading papers are key skills for scientists. Indeed, success at publishing is used to evaluate scientists [1] and can help predict their future success [2]. In the production and consumption of papers, multiple parties are involved, each having their own motivations and priorities. The editors want to make sure that the paper is significant, and the reviewers want to determine whether the conclusions are justified by the results. The reader wants to quickly understand the conceptual conclusions of the paper before deciding whether to dig into the details, and the writer wants to convey the important contributions to the broadest audience possible while convincing the specialist that the findings are credible. You can facilitate all of these goals by structuring the paper well at multiple scales-- spanning the sentence, paragraph, section, and document.

Clear communication is also crucial for the broader scientific enterprise because ‘concept transfer’ is a rate-limiting step in scientific cross-pollination. This is particularly true in the biological sciences and other fields that comprise a vast web of highly interconnected sub-disciplines. As scientists become increasingly specialized, it becomes more important (and difficult) to strengthen the conceptual links. Communication across disciplinary boundaries can only work when manuscripts are readable, credible, and memorable.

The claim that gives significance to your work has to be supported by data and a logic that gives it credibility. Without carefully planning the paper’s logic, there will often be missing data or missing logical steps on the way to the conclusion.

While these lapses are beyond our scope, your scientific *logic* must be crystal clear to powerfully make your *claim*.

Here we present 10 simple rules for structuring papers (see Table 1). The first four rules are *principles* that apply to all the parts of a paper and further, to other forms of communication such as grants and posters. The next four rules deal with the primary goals of each of the main *parts* of papers. The final two rules deliver guidance on the *process* – heuristics for efficiently constructing manuscripts.

**Table 1:** A summary of the 10 rules and how to tell if they are being violated

|  |  |
| --- | --- |
| **Rule** | **Sign it is violated** |
| 1: One big idea | Readers cannot give one-sentence summary |
| 2: Humans as audience | Readers do not ‘get’ the paper |
| 3: Context, Content, Conclusion | Readers ask why something matters or what it means |
| 4: Optimize logical flow | Readers stumble on a small section of the text |
| 5: Abstract: Compact summary of paper | Readers cannot give the ‘elevator pitch’ of your work after reading it |
| 6: Introduction: Why the paper matters | Readers show little interest in the paper |
| 7: Results: Why the conclusion is justified | Readers do not agree with your conclusion |
| 8: Discussion: Preempt criticism, give future impact | Readers are left with unanswered criticisms/questions on their mind |
| 9: Time allocation | Readers struggle to understand your central contribution despite your having worked hard. |
| 10: Iterate the story | The paper’s contribution is rejected by test readers, editors, or reviewers. |

## Principles (rules 1-4)

Writing is communication. Thus, the reader’s experience is of primary importance, and all writing serves this goal. When you write, you should constantly have your reader in mind. These four rules help you to avoid losing your reader.

### Rule 1: Focus your paper on one central contribution, which you communicate in the title

Your communication efforts are successful if readers can still describe the main contribution of your paper to their colleagues a year after reading it. While it is clear that a paper often needs to communicate a number of innovations on its way to its final message, *it does not pay to be greedy*. Focus on a single message: papers that simultaneously focus on multiple contributions tend to be less convincing about each and are therefore less memorable.

The most important element of a paper is the title-- think of the ratio of the number of titles you read to the number of papers you read. The title is typically the first element a reader encounters, so its quality [3] determines whether the reader will invest time in reading the abstract.

The title not only transmits the paper’s central contribution, but can also serve as a constant reminder (to you) to focus the text on transmitting that idea. Science is, after all, the abstraction of simple principles from complex data. The title is the ultimate refinement of the paper’s contribution. Thinking about the title early--and regularly returning to hone it--can help not only the writing of the paper, but also the process of designing experiments or developing theories.

This Rule of One is the most difficult rule to optimally implement, because it comes face to face with the key challenge of science: making the claim/model as simple as data and logic can support, but no simpler. In the end, your struggle to find this balance may appropriately result in “one contribution” that is multifaceted. For example, a technology paper may describe both its new technology and a biological result using it; the bridge that unifies these two facets is a clear description of how the new technology can be used to do new biology.

### Rule 2: Write for flesh-and-blood human beings who do not know your work

Because you are the world’s leading expert at exactly what you are doing, you are also the world’s least qualified person to judge your writing from the perspective of the naïve reader. The majority of writing mistakes stem from this predicament. Think like a designer—for each element, determine the impact that you want to have on people and then strive to achieve that objective [4]. Try to think through the paper like a naïve reader who must first be made to care about the problem you are addressing (see rule 6), and then wants to understand your answer with minimal effort.

Define technical terms clearly, because readers can become frustrated when they encounter a word they don’t understand. Avoid abbreviations and acronyms, so that readers do not have to go back to earlier sections to identify them.

The vast knowledge base of human psychology is useful in paper writing. For example, people have working-memory constraints: they can only remember a small number of items and are better at remembering the beginning and the end of a list than the middle [5]. Do your best to minimize the number of loose threads that the reader has to keep in mind at any one time.

### Rule 3: Stick to the context-content-conclusion (C-C-C) scheme

The vast majority of popular (i.e., memorable and re-tellable) stories have a structure with a discernible beginning, a well-defined body, and an end. The beginning sets up the context for the story, while the body (content) advances the story towards an ending where the problems find their conclusion. This structure reduces the chances that the reader will wonder “why was I told that?” (if the context is missing) or “so what?” (if the conclusion is missing).

There are many ways of telling a story. Mostly, they differ in how well they serve a patient reader versus an impatient one [6]. The impatient reader needs to be engaged quickly-- this can be accomplished by presenting the most exciting content first (e.g., as seen in news articles). The C-C-C scheme that we advocate serves a more patient reader, who is willing to spend the time to get oriented with the context. A consequent disadvantage of C-C-C is that it may not optimally engage the impatient reader. This disadvantage is mitigated by the fact that the structure of scientific articles, specifically the primacy of the title and abstract, already forces the content to be revealed quickly. Thus, a reader who proceeds to the Introduction is likely engaged enough to have the patience to absorb the context. Further, one hazard of excessive ‘content first’ story structures in science is that you may generate skepticism in the reader, since they may be missing an important piece of context that makes your claim more credible. For these reasons, we advocate C-C-C as a ‘default’ scientific story structure.

The C-C-C scheme defines the structure of the paper on multiple scales. At the whole-paper scale, the introduction sets the context, the results are the content, and the discussion brings home the conclusion. Applying C-C-C at the paragraph scale, the first sentence defines the topic or context, the body hosts the novel content put forth for the reader’s consideration, while the last sentence provides the conclusion to be remembered.

Deviating from the C-C-C structure often leads to papers that are hard to read, but writers often do so because of their own autobiographical context. During our everyday life as scientists, we spend a majority of our time producing content and a minority amidst a flurry of other activities. We run experiments, develop the exposition of available literature, and combine thoughts using the magic of human cognition. It is natural to want to record these efforts on paper and structure a paper chronologically. But for our readers, most details of our activities are extraneous. They do not care about the chronological path by which you reached a result; they just care about the ultimate claim and the logic supporting it (see rule 7). Thus, all our work must be reformatted to provide a context that makes our material meaningful and a conclusion that helps the reader to understand and remember it.

### Rule 4: Optimize your logical flow by avoiding zig-zag and using parallelism

#### Avoiding zig-zag

Only the central idea of the paper should be touched upon multiple times. Otherwise each subject should be covered in only one place to minimize the number of subject changes. Related sentences or paragraphs should be strung together rather than being interrupted by unrelated material. Ideas that are similar, such as two reasons why we should believe something, should come one immediately after the other.

#### Using parallelism

Similarly, across consecutive paragraphs or sentences, parallel messages should be communicated with parallel form. Parallelism makes it easier to read the text because the reader is familiar with the structure. For example, if we have three independent reasons why we prefer one interpretation of a result over another, it is helpful to communicate them with the same syntax so that this syntax becomes transparent to the reader, allowing them to focus on the content. There is nothing wrong with using the same word multiple times in a sentence or paragraph. Resist the temptation to use a different word to refer to the same concept—this makes readers wonder if the 2^nd^ word has a differently nuanced meaning.

## The components of a paper (Rules 5-8)

The individual parts of a paper—abstract, introduction, results, and discussion— have different objectives and thus they each apply the C-C-C structure a little differently in order to achieve their objectives. We will discuss these specialized structures in this section, and summarize them in Figure 1.

**Figure 1:**
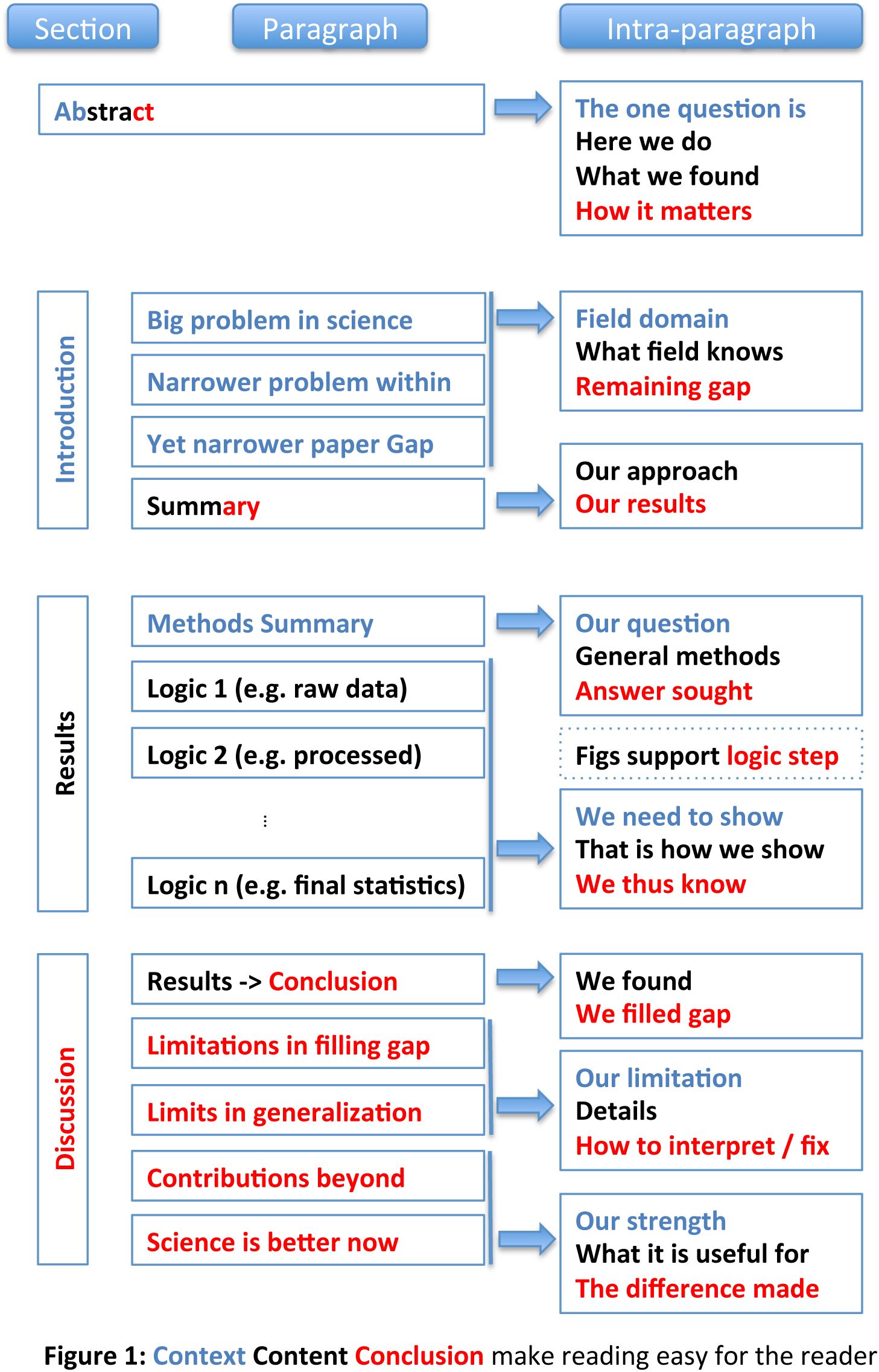
Summary of a paper’s structural elements at three spatial scales: across sections, across paragraphs, and within paragraphs. Note that the abstract is special in that it contains all 3 elements (Context, Content, and Conclusion), thus comprising all 3 colors.

### Rule 5: Tell a complete story in the abstract

The abstract is, for most readers, the only part of the paper that will be read. This means that the abstract must convey the entire message of the paper effectively. To serve this purpose, the abstract’s structure is highly conserved. Each of the C-C-C elements is detailed below.

The *context* must get across the gap that the paper will fill. The first sentence orients the reader by introducing the broader field in which the particular research is situated. Then this context is narrowed until it lands on the open question that the research answered. A successful context section sets the stage for distinguishing the paper’s contributions from the current state of the art by communicating what is missing in the literature (i.e., the specific gap) and why that matters (i.e., the connection between the specific gap and the broader context that the paper opened with).

The *content* (“Here we”) first describes the novel method or approach that you used to fill the gap/question. Then you present the meat—your executive summary of the results.

Finally, the *conclusion* interprets the results to answer the question that was posed at the end of the context section. There is often a 2^nd^ part to the conclusion section, which highlights how this conclusion moves the broader field forward (i.e., ‘broader significance’). This is particularly true for more ‘general’ journals with a broad readership.

This structure helps you avoid the most common abstract mistake: talking about results before the reader is ready to understand them. Good abstracts usually take many iterations of refinement to make sure the results fill the gap like a key fits its lock. The broad-narrow-broad structure allows you to communicate with a wider readership (through breadth) while maintaining the credibility of your claim (which is always based on a finite/narrow set of results).

### Rule 6: Get across why the paper matters in the introduction

The introduction highlights the gap that exists in current knowledge or methods, and why it is important. This is usually done by a set of progressively more specific paragraphs that culminate in a clear exposition of what is lacking in the literature, followed by a paragraph summarizing what the paper does to fill that gap.

As an example of the progression of gaps, a first paragraph may explain why understanding cell differentiation is an important topic and that the field has not yet solved what triggers it (a field gap). A second paragraph may explain what is unknown about the differentiation of a specific cell type, such as astrocytes (a subfield gap). A third may provide clues that a particular gene might drive astrocytic differentiation, and then state that this hypothesis is untested (the gap within the subfield that you will fill). The gap statement sets the reader’s expectation for what the paper will deliver.

The structure of each Introduction paragraph (except the last) serves the goal of developing the gap. Each paragraph first orients the reader to the topic (a context sentence or two) and then explains the ‘knowns’ in the relevant literature (content) before landing on the critical ‘unknown’ (conclusion) that makes the paper matter at the relevant scale. Along the path, there are often clues given about the mystery behind the gaps; these clues lead to the untested hypothesis or undeveloped method of the paper and give the reader hope that the mystery is solvable. The introduction should not contain a broad literature review beyond the motivation of the paper. This gap-focused structure makes it easy for experienced readers to evaluate the potential importance of a paper – they just need to assess the importance of the claimed gap.

The last paragraph of the introduction is special – it compactly summarizes the results, which fill the gap you just established. It differs from the abstract in several ways: it does not need to present context (which has just been given above), it is somewhat more specific about the results, and it only briefly previews the conclusion of the paper, if at all.

### Rule 7: Communicate the results as a sequence of statements, supported by figures, that connect logically to support the central contribution

The results section needs to convince the reader that the central claim is supported by data and logic. Every scientific argument has its own particular logic structure, which dictates the sequence in which its elements should be presented.

For example, one paper may set up a hypothesis, verify that a method for measurement is valid in the system under study, and then use the measurement to disprove the hypothesis. Alternatively, a paper may set up multiple alternative (and mutually exclusive) hypotheses, and then disprove all but one to provide evidence for the remaining interpretation. The fabric of the argument will contain controls and methods where they are needed for the overall logic.

In the outlining phase of paper preparation (see rule 9), sketch out the logic structure of how your results support your claim and convert this into a sequence of declarative statements that become the headers of subsections within the results section (and/or the titles of figures). Most journals allow this type of formatting, but if your chosen journal does not, these headers are still useful during the writing phase and can be either adapted to serve as introductory sentences to your paragraphs or deleted before submission. Such a clear progression of logical steps makes the paper easy to follow.

Figures, their titles, and legends are particularly important because they show the most objective support (data) of the steps that culminate in the paper’s claim. Moreover, figures are often viewed by readers who skip directly from the abstract in order to save time. Thus the title of the figure should communicate the conclusion of the analysis, and the legend tells how it was done. Figure-making is an art unto itself; the Edward Tufte books remain the gold standard for learning this craft [7,8].

The first results paragraph is special in that it typically summarizes the overall approach to the problem outlined in the Introduction, along with any key innovative methods that were developed. Most readers do not read the methods, so this paragraph gives them the gist of the methods that were used.

Each subsequent paragraph in the results section starts with a sentence or two, setting up the question that the paragraph answers. For example “To verify that there are no artifacts,…“, “What is the test-retest reliability of our measure?“, or “We next tested whether Ca^2+^ flux through L-type Ca^2+^ channels was involved“. The middle of the paragraph presents data and logic that pertain to the question, and the paragraph ends with a sentence that answers the question. For example, it may conclude that none of the potential artifacts was detected. This structure makes it easy for experienced readers to fact-check a paper. Each paragraph convinces the reader of the answer given in its last sentence. This makes it easy to find the paragraph where a suspicious conclusion is drawn, and check the logic of that paragraph. The result of each paragraph is a logical statement, and paragraphs farther down in the text rely on the logical conclusions of previous paragraphs, much as theorems are built in mathematical literature.

### Rule 8: Discuss how the gap was filled, the limitations of the interpretation, and the relevance to the field

The discussion section explains how the results have filled the gap identified in the introduction, provides caveats to the interpretation, and describes how the paper advances the field by providing new opportunities. This is typically done by recapitulating the results, discussing the limitations, and then revealing how the central contribution may catalyze future progress. The first discussion paragraph is special in that it generally summarizes the important findings from the results section. Some readers skip over substantial parts of the results, so this paragraph at least gives them the gist of that section.

Each following paragraph in the discussion section starts by describing an area of weakness or strength of the paper. It then evaluates the strength or weakness by linking it to the relevant literature. Discussion paragraphs often conclude by describing a clever, informal way of perceiving the contribution or by discussing future directions that can extend the contribution.

For example, the first paragraph may summarize the results, focusing on their meaning. The second through fourth paragraphs may deal with potential weaknesses and how the literature alleviates those concerns or how future experiments can deal with these weaknesses. The fifth paragraph may then culminate in a description of how the paper moves the field forward. Step by step, the reader thus learns to put the paper’s conclusions into the right context.

## Process (Rules 9-10)

To produce a good paper, authors can use helpful processes and habits. Some aspects of a paper affect its impact more than others, suggesting that your investment of time should be weighted towards the issues that matter most. Moreover, iteratively using feedback from colleagues allows the authors to improve the story at all levels to produce a powerful manuscript. Choosing the right process makes writing papers easier and more effective.

### Rule 9: Allocate time where it matters: Title, abstract, figures, and outlining

The central logic that underlies a scientific claim is paramount. It is also the bridge connecting the experimental phase of a research effort with the paper-writing phase. Thus, it is useful to formalize the logic of ongoing experimental efforts (e.g., during lab meetings) into an evolving document of some sort that will ultimately steer the outline of the paper.

You should also allocate your time according to the importance of each section. The title, abstract, and figures are viewed by far more people than the rest of the paper, and the methods section is read least of all. Budget accordingly.

The time that we do spend on each section can be used efficiently by planning text before producing it. Make an outline. We like to write one informal sentence for each planned paragraph. It is often useful to start the process around descriptions of each result – these may become the section headers in the results section. As the story has an overall arc, each paragraph should have a defined role in advancing this story. This role is best scrutinized at the outline stage, to reduce wasting time on wordsmithing paragraphs that don’t end up fitting within the overall story.

### Rule 10: Get feedback to reduce, reuse, and recycle the story

Writing can be considered an optimization problem in which you simultaneously improve the story, the outline, and all the component sentences. In this context, it is important not to get too attached to one’s writing. In many cases, trashing entire paragraphs and rewriting is a faster way to produce good text than incremental editing.

There are multiple signs that further work is necessary on a manuscript (see Table 1). For example, if you, as the writer, cannot describe the entire outline of a paper to a colleague in a few minutes, then clearly a reader will not be able to. You need to further distill your story. Finding such violations of good writing helps to improve the paper at all levels.

Successfully writing a paper typically requires input from multiple people. Test readers are necessary to make sure that the overall story works. They can also give valuable input on where the story appears to move too quickly or too slowly. They can clarify when it is best to go back to the drawing board and retell the entire story. Reviewers are also extremely useful. Non-specific feedback and unenthusiastic reviews often imply that the reviewers did not ‘get’ the big-picture storyline. Very specific feedback usually points out places where the logic within a paragraph was not sufficient. It is vital to accept this feedback in a positive way. As input from others is essential, a network of helpful colleagues is fundamental to making a story memorable. To keep this network working, make sure to pay back your colleagues by reading their manuscripts.

## Discussion

This paper focused on the structure or ‘anatomy’ of manuscripts. We had to gloss over many finer points of writing, including word choice and grammar, the creative process, and collaboration. A paper about writing can never be complete; as such there is a large literature dealing with issues of scientific writing [9,10,11,12,13,14,15,16,17].

Personal style often leads writers to deviate from a rigid, conserved structure and it can be a delight to read a paper that creatively bends the rules. However, as with many other things in life, a thorough mastery of the standard rules is necessary to successfully bend them [18]. In following these guidelines, scientists will be able to address a broad audience, bridge disciplines, and more effectively enable integrative science.

